# A balanced circadian glycolytic rhythm drives cardiomyocyte cell cycle progression during fish heart regeneration

**DOI:** 10.1101/2025.09.15.675600

**Authors:** Gennaro Ruggiero, Rita Alonaizan, William T. Stockdale, Konstantinos Lekkos, Denise Posadas Pena, Vanessa Obi, Jun Ying, Sara Al-Dahwi, John Walsby-Tickle, Madeleine E. Lemieux, Gilbert Weidinger, Mathilda T.M. Mommersteeg

## Abstract

The ability of heart tissue to repair itself after injury has fascinated scientists for decades^1,2^. Researchers have long studied the internal body clock, or circadian rhythm, for its role in coordinating daily cycles of metabolism and cell activity^3,4^, but its relevance to heart repair has remained unknown. This study explores, for the first time, whether natural daily rhythms influence heart regeneration—a process driven by cardiomyocyte proliferation. We discovered that DNA replication, mitosis, oxidative phosphorylation, and glycolysis follow a precise daily order in regenerating zebrafish hearts. Disrupting core clock gene expression abolishes the rhythms of glycolysis and mitosis, preventing cardiomyocyte cell cycle progression and regeneration. Insulin-resistant *Astyanax mexicanus* cavefish, which have adapted to dark caves, similarly show a loss of mitosis rhythm and cardiomyocyte cell cycle progression, which we find is caused by reduced glycolysis. Despite this reduction, glycolysis rhythm displays a larger amplitude in cavefish—a pattern recapitulated in insulin-resistant zebrafish. Insulin resistance resets metabolic rhythms to the morning, which is equally detrimental to regeneration. Here, we show that successful cardiac regeneration depends on synchronised clock and glucose rhythms, which together orchestrate the cell cycle events essential for cardiomyocyte proliferation and tissue repair.

## Main

Cardiovascular disease is the leading cause of death worldwide due in part to the limited capacity of the adult mammalian heart to regenerate ^5^. Accumulating evidence indicates that cell-autonomous timekeeping mechanisms, termed circadian clocks, adjust cardiovascular functions to a 24-hour cycle, allowing the synchronisation of cellular cardiac processes with diurnal rhythms in the environment ^6–16^. This is highlighted by increased incidence of myocardial infarction (MI) onset in the early morning ^17–19^, and by the observed correlation between infarct size and the time of day at which MI occurs ^20–22^. Daily oscillation of biological functions has been observed in various cardiovascular cells, including endothelial cells ^23^, vascular smooth muscle cells ^24^, and cardiomyocytes ^11,25–27^. Also, the regenerative process has been shown to be intricately regulated by the biological clock across a diverse array of cell types and organs such as the skin, where circadian rhythms impact wound healing and cell turnover ^28–31^, the intestine, where they govern epithelial renewal and barrier maintenance ^32–34^, and the liver, where daily cycles control hepatocyte proliferation ^35–39^. These findings highlight the universal role of circadian clocks in synchronising regenerative processes with physiological and environmental rhythms. However, whether the circadian clock modulates heart regeneration in vertebrates remains unknown. Here we investigated the interplay between the circadian clock and cardiac regeneration, using *Danio rerio* (zebrafish) and *Astyanax mexicanus* as model organisms, well-established species for studying heart regeneration ^1,40^.

### The cardiomyocyte cell cycle is rhythmic during heart regeneration

Circadian control of gene expression is regulated by a transcription-translation feedback loop of core clock genes (Extended Data Fig. 1a). After verifying rhythmicity of core clock gene expression in the uninjured zebrafish heart and that this rhythm was maintained after injury (Extended Data Fig. 1b-h), we performed bulk RNA sequencing (RNAseq) to identify regeneration-dependent changes in circadian gene expression. Hearts of wild-type (WT) zebrafish were cryoinjured, and ventricles collected at 4-hour intervals over a total of 48 hours, starting at 7 days post cryoinjury (dpci), as well as uninjured controls (Fig. 1a). Analysis using the *dryR* algorithm ^41^ showed that ∼12% of genes in the heart were rhythmic, either only before or after injury, or at any state. Analysis for processes specifically becoming rhythmic after injury revealed that this subset of 322 clock-controlled genes (CCGs) was strongly enriched in functions linked to the cell cycle and in particular mitosis (Fig. 1b-c). The top enriched term *mitotic cell cycle process* with genes as *aurkb*, *ect2*, and *cdk1* was highly rhythmic at 7dpci, the height of cardiomyocyte proliferation, with the peak of expression at the end of the light period (Fig. 1d and Extended Data Fig. 1i). While the top enriched terms highlighted the rhythmicity of the M-phase, also genes involved in S-phase were rhythmic specifically after injury, including *pcna* (Fig. 1e).

**Figure 1.**
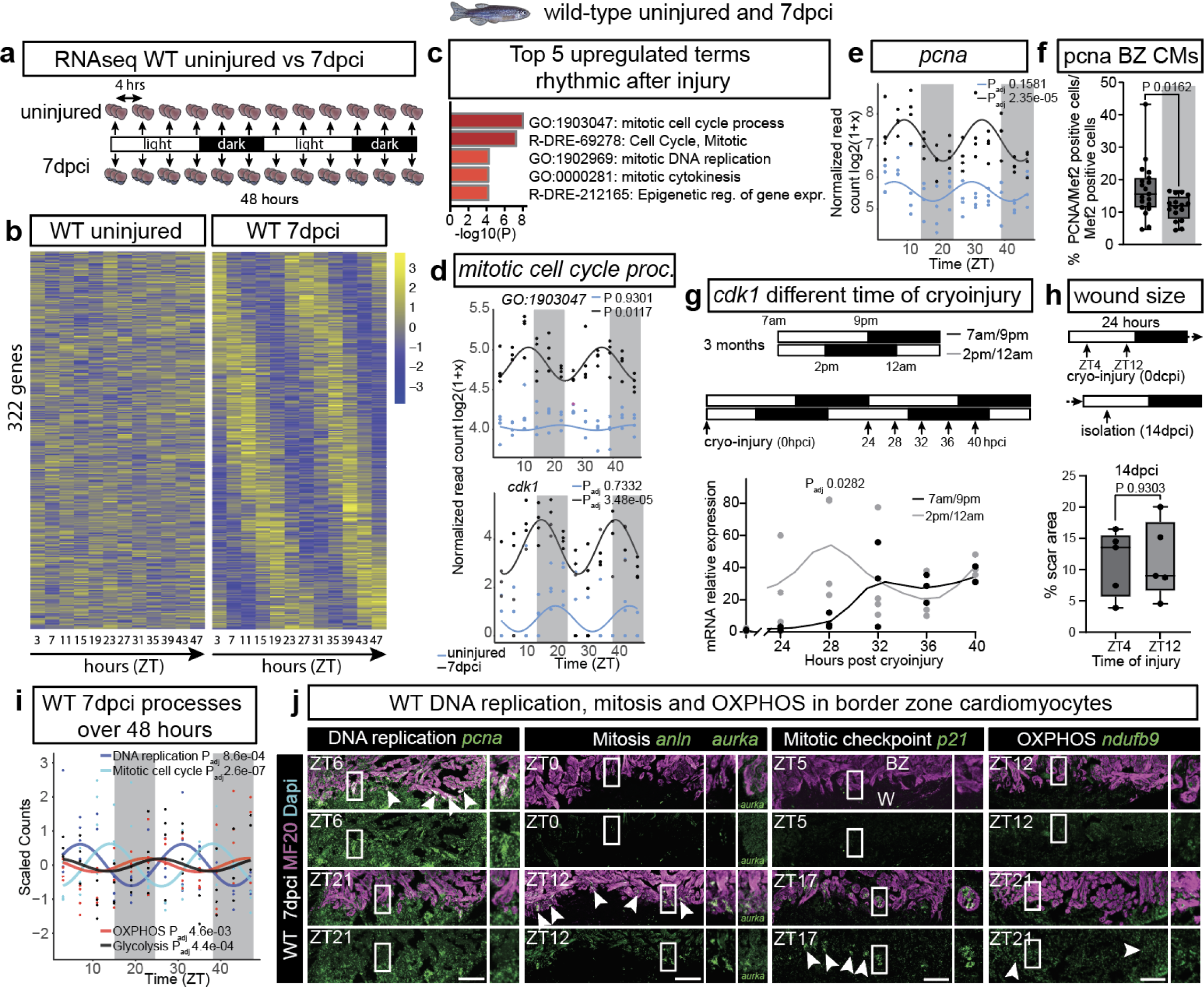
The cardiomyocyte cell cycle is rhythmic during heart regeneration. **a**, RNAseq experimental design schematic. Horizontal bars indicate light (14h) and dark (10h) periods, before and during sampling of uninjured and 7 dpci WT hearts. **b**, Heatmap showing rhythmic gene expression gain post-injury for 322 CCGs identified by *dryR*. Genes are organised by peak expression timing (ZT), with each column representing a single time point (biological replicates ≥ 4 sampled every 4h over 48h). **c**, Bar graph showing Metascape-enriched terms ^42^ for the 322 rhythmic genes post-injury, coloured by p-values. **d**,**e**, Rhythmic expression of genes within the GO term *mitotic cell cycle process* and *cdk1* (**d**) and *pcna* (**e**) in uninjured vs. 7 dpci WT hearts. **f**, PCNA/Mef2-positive BZ CMs quantification by immunofluorescence in 7 dpci WT hearts between light and dark periods (unpaired two-tailed Welsh t-test). **g**, WT zebrafish were kept in control (7am–9pm light) or shifted (12am–2pm light) cycles for 3 months. Cryoinjury was performed at the same time which corresponded to ZT0 in control or ZT7 in shifted cycles. Hearts were collected every 4 hours from 24 to 40 hours post-cryoinjury (hpci) for *cdk1* qRT-PCR analysis (two-way ANOVA with Sidak). **h**, Scar area quantification at 14 dpci in WT hearts injured at ZT4 vs. ZT12 (unpaired two-tailed Student’s t-test). **i**, Rhythmic expression of *DNA replication*, *Mitotic cell cycle*, *OXPHOS*, and *Glycolysis* in 7 dpci WT hearts. **j**, RNAscope analysis of *pcna*, *anln*, *aurka*, *p21*, and *ndufb9* in 7 dpci WT heart sections, co-stained with MF20. (**e**, **d**, **i**) *dryR* analysis. Scale bars 100 µm. BZ, border zone cardiomyocytes; W, wound.

Since immunohistochemistry for PCNA in border-zone cardiomyocytes (BZ CMs) is widely used in the field as an indicator of regeneration, we analysed the number of PCNA-positive BZ CMs during the day versus night. We observed a higher number of PCNA-positive BZ CMs during the light period, confirming rhythmic expression at protein level (Fig. 1f). Shifting the start of the light/dark cycle by 7 hours showed that the initiation of cardiac cell proliferation after injury was governed by the phase of the light/dark cycle rather than the timing of the injury, as shown by *cdk1* gene expression (Fig. 1g). Cryoinjury in the morning versus the evening did not influence scar size at 14 dpci, indicating that the timing of injury did not influence long-term regeneration (Fig. 1h). Together, these findings demonstrate that critical processes for heart regeneration, including the cell cycle, are highly rhythmic in BZ CMs. We then divided the 24-hour circadian RNAseq data at 7dpci into blocks of 4 hours, which showed that DNA replication occurred predominantly in the morning (Zeitgeber Time, ZT4-7), approximately 8 hours before the peak in mitosis (ZT12-15). This was followed by a peak in the expression of genes involved in oxidative phosphorylation (OXPHOS) at night (ZT20-23), while glycolysis peaked just after lights switched on (ZT0-4) (Extended Data Fig. 1j and Fig. 1i). Analysis of sections confirmed that these rhythmic processes were localised in the BZ CMs, adjacent to the injury site (Fig. 1j).

### Disrupting circadian rhythm inhibits heart regeneration and alters the myocardial glycolytic response

To establish if clock-regulated rhythmicity is important for regeneration, we analysed a zebrafish line in which three core clock genes, *cry1a*, *cry3a*, and *per2*, were mutated (referred to as TKO, triple knock-out) ^43^. The TKO indeed showed larger wounds compared to WT controls at both 7 and 21 dpci (Extended Data Fig. 2a and Fig. 2a), with reduced levels of cardiomyocyte proliferation at 7 dpci (Fig. 2b), indicating that the core clock machinery is important for regeneration. Bulk RNAseq on TKO and WT control ventricles, isolated every 4 hours over 24 hours at 7 dpci (Fig. 2c) showed that while core clock genes were largely still rhythmically expressed in the TKO, their phase, amplitude, and expression level were changed compared to WTs, highlighted for *cry1a*, and *bmal1b*, and *per1b* (Extended Data Fig. 2b-d), with an overall phase advance shift (Fig. 2d). While rhythm was not completely lost in the TKO, a shift in rhythms is sufficient to disrupt cellular processes, as also highlighted by susceptibility to disease in shift workers ^44^. As a result of this shift, many genes completely lost their rhythmic expression, with *cell cycle* and *cell division* emerging as top affected processes in the TKO (Fig. 2e). While DNA replication peaked in the morning, similar to the WT, the rhythm of genes involved in later stages of the cell cycle was dampened in the TKO compared to the WT (Fig. 2f-h).

**Figure 2.**
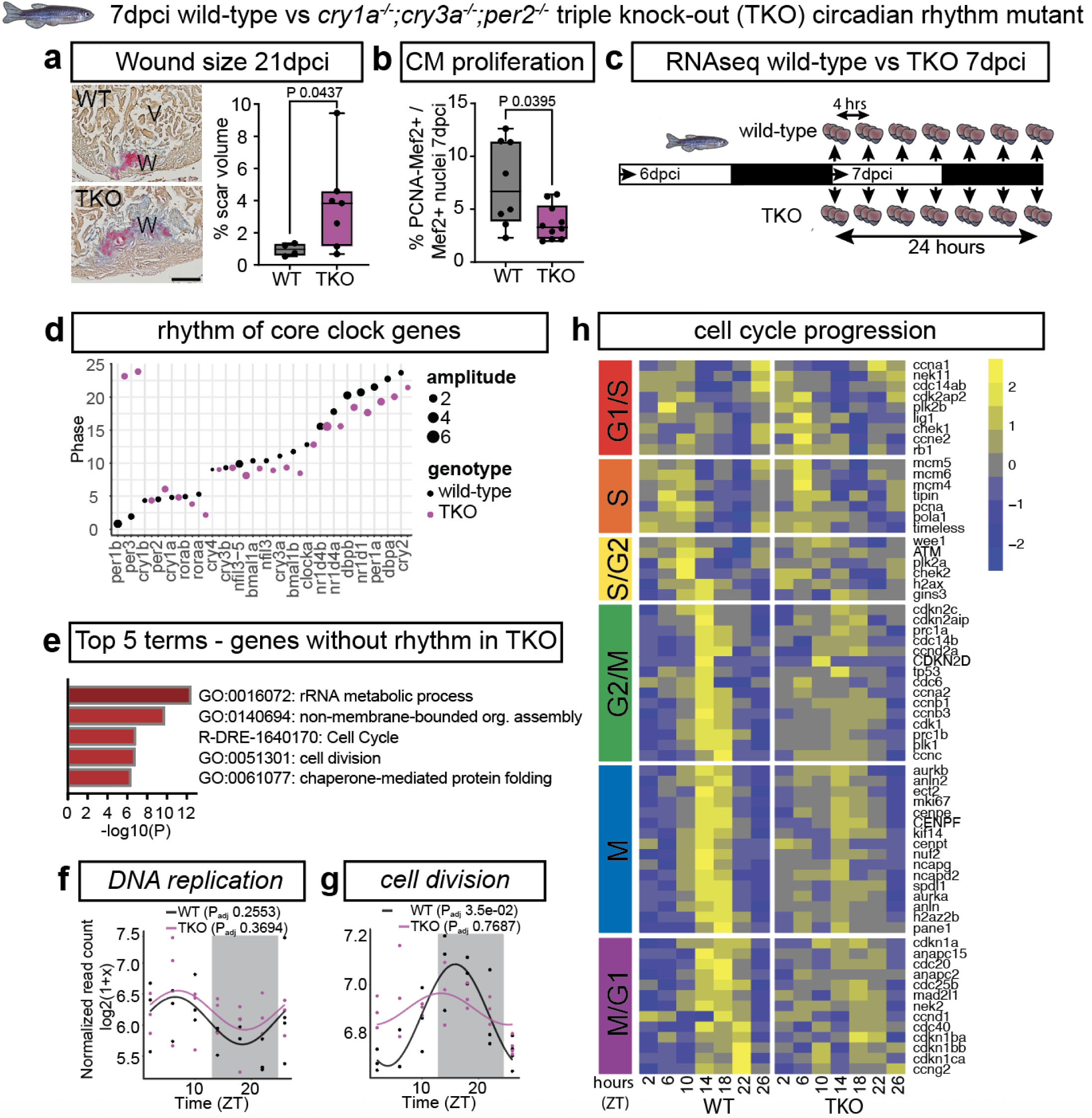
Disrupting circadian rhythm inhibits heart regeneration. **a**, AFOG staining showing collagen (blue) and fibrin (red) scars in TKO vs. WT at 21 dpci. Scar volume quantification (unpaired two-tailed Welch t-test). **b**, PCNA/Mef2-positive BZ CMs quantification in WT vs. TKO hearts at 7 dpci (unpaired two-tailed Welch t-test). **c**, RNAseq experimental design schematic. Horizontal bars indicate light (12h) and dark (12h) periods before and during sampling of 7 dpci WT and TKO hearts. **d,** Dot plot showing phase and amplitude differences of core clock genes in WT vs. TKO hearts at 7 dpci. **e**, Bar graph showing Metascape-enriched terms for genes that lost rhythm in TKO hearts at 7 dpci, coloured by p-values. **f**,**g**, Rhythmic expression profiles of *DNA replication* and *cell division* in WT and TKO hearts at 7 dpci. **h**, Heatmap of cell cycle rhythmic genes in WT and TKO hearts at 7 dpci. Each column represents a time point, (biological replicates ≥ 3 sampled at 4-hour intervals over 24 hours). (**d**, **f** to **h**) *dryR* analysis. Scale bars 100 µm. V, ventricle; W, wound.

This suggested that proliferating cells in the TKO initiate DNA replication but might not pass cell cycle checkpoints, blocking progression towards M-phase. We confirmed this by measuring DNA content of BZ CMs, showing lack of progression from S- into M-phase (Fig. 3a). Replication stress and DNA damage are major causes of cell cycle stalling ^45,46^, and excessive replication stress inhibits zebrafish heart regeneration ^47^. Indeed, we observed upregulation of DNA damage checkpoint genes such as *chek1* (Fig. 2h) and increased levels of γH2a.x in TKO BZ CMs indicating cell cycle delay (Fig 3b).

**Figure 3.**
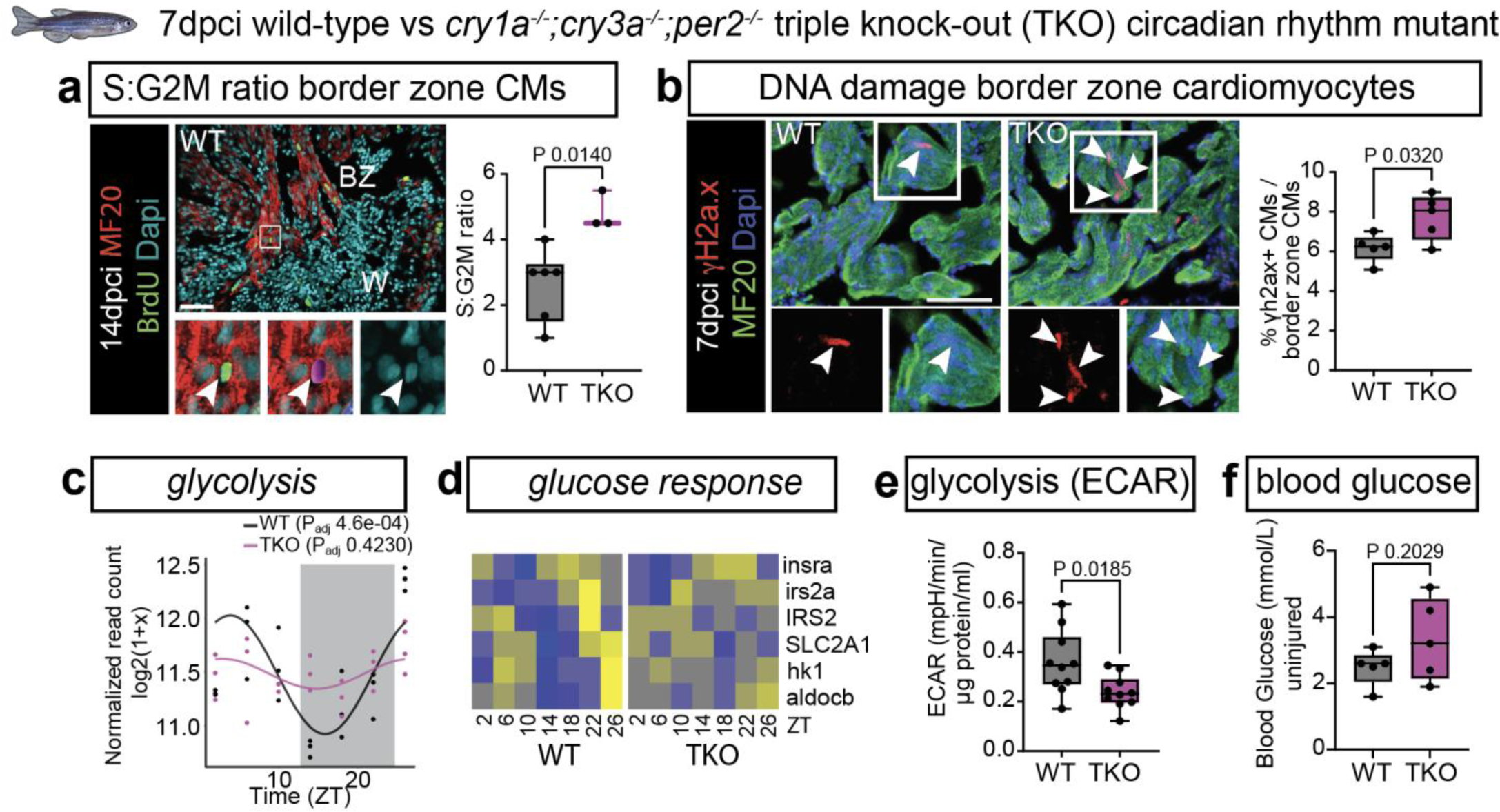
Disrupting circadian rhythm alters the myocardial glycolytic response. **a**, WT and TKO zebrafish were injected with BrdU at 7 dpci and hearts were isolated at 14 dpci. Image of BZ CM DNA content analysis, highlighting G1/2n, S, and G2M/4n nuclei, stained with BrdU, Mef2, and DAPI. S:G2M ratio comparison of BZ CM between WT vs. TKO at 14 dpci (unpaired two-tailed Student’s t-test), **b,** γH2a.x/MF20-positive BZ CMs quantification in WT vs. TKO hearts at 7 dpci (unpaired two-tailed Student’s t-test). **c**, Rhythmic expression of *glycolysis* in WT and TKO hearts at 7 dpci. **d**, Heatmap of glucose response rhythmic genes in WT vs. TKO at 7 dpci. **e**, Glycolysis - Extracellular acidification rate (ECAR) in WT vs. TKO hearts at 7 dpci (unpaired two-tailed Student’s t-test). **f**, Fasting blood glucose levels in WT and TKO (unpaired two-tailed Student’s t-test). (**c** and **d**) *dryR* analysis. Scale bars 100 µm. BZ, border zone cardiomyocytes; V, ventricle; W, wound.

Cardiomyocytes rely on glycolysis for proliferation ^48,49^. As glycolysis is clock controlled ^50^, we analysed the rhythms of genes involved in glycolysis. These genes had completely lost their rhythm in the TKO (Fig. 3c), in particular genes involved in insulin signalling (*insra*, *irs2a*, *IRS2*), glucose transport into the cell (*SLC2A1,* the main glucose transporter in the fish heart ^51^) and the initial steps of glycolysis (*hk1*, *aldocb*) (Fig. 3d). This resulted in functionally lower levels of glycolysis in TKO hearts (Fig. 3e), despite normal feeding and comparable blood glucose levels between WT and TKO (Fig. 3f). This data shows that disturbing the core clock mechanism abolishes the rhythms of both cell division and glycolysis, inhibiting regeneration. Although the core clock may directly regulate cell cycle progression ^52^, glycolysis-derived energy and nucleotides are essential for replication fork progression and BZ CM proliferation ^49^, making the defective glycolysis rhythm a likely cause of the cell cycle stalling.

### Reduced glycolysis is responsible for the stalling of cardiomyocyte cell cycle progression

We have previously shown that *Astyanax mexicanus* surface river fish can regenerate their hearts, while blind cavefish from the Pachón population cannot and instead form a permanent scar ^40^. Adaptation to perpetual darkness in the cave environment led to the disruption of the endogenous circadian clock ^53^. As our results show that circadian rhythm is key to successful regeneration, we questioned if rhythmic processes and cell cycle regulation were also affected in Pachón cavefish, as this could provide further insight into how circadian rhythm affects BZ CM cell cycle progression. Therefore, we performed bulk RNAseq at 4-hour intervals over 24 hours on both surface and Pachón hearts at 7 dpci (Fig. 4a). Animals were kept in the dark during these 24 hours to distinguish between light-inducible mechanisms and those purely governed by the internal circadian clock (circadian time (CT)). The results revealed significant phase delay of the core clock genes between the two morphs (Fig. 4b), contrasting the phase advance observed in the TKO. Many genes lost their rhythmicity in Pachón, although surprisingly, other genes seemed rhythmic specifically in Pachón (Fig. 4c,d). Despite the contrasting phase shift to the TKO, mitosis rhythm was completely absent in Pachón, including that of the key cell cycle regulator *cdk1*, while mitosis was the dominant process in surface fish at CT 16-19 (Fig. 4e-g).

**Figure 4:**
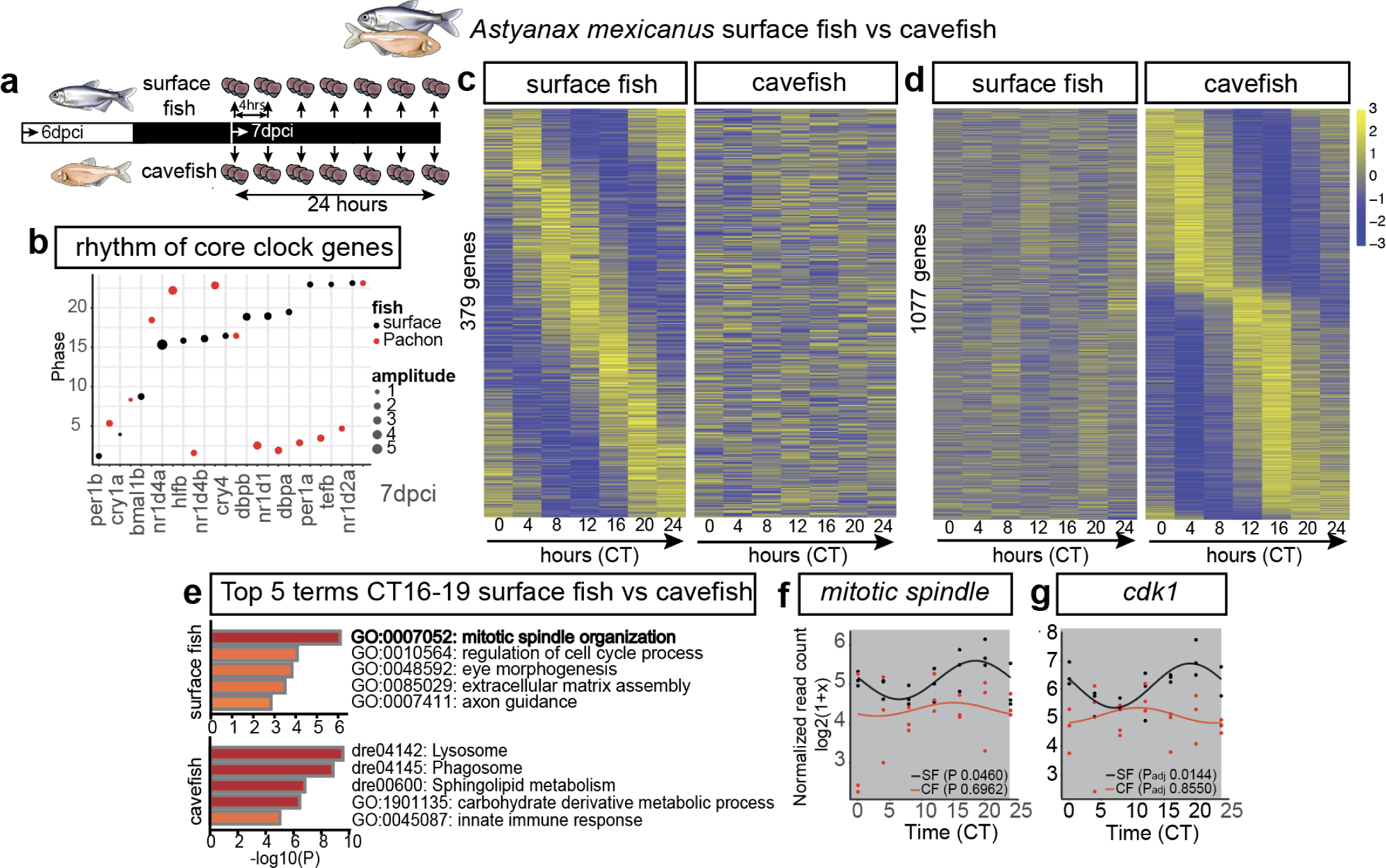
Circadian clock gene expression varies during heart regeneration in *Astyanax mexicanus*. **a**, RNAseq experimental design schematic. Horizontal bars indicate light (14h) and dark (10h) periods before sampling of Pachón and surface fish hearts at 7 dpci. All sampling was conducted in complete darkness. **b**, Dot plot showing phase and amplitude differences in core clock genes between Pachón and surface hearts at 7 dpci. **c**,**d**, Heatmap of genes identified as rhythmic in Pachón or surface hearts at 7 dpci (biological replicates ≥ 3 sampled at 4-hour intervals over 24 hours). **e**, Bar graph of Metascape-enriched terms for CCGs in surface hearts at 7 dpci. Genes peak within the 16–19 Circadian Time (CT) window, coloured by p-values. **f**, **g**, Rhythmic expression of the genes representing *mitotic spindle* process and *pcna* in Pachón and surface hearts at 7dpci. (**b** to **d**, **f, g**) *dryR* analysis.

Similar to the TKO, this resulted in lack of cell cycle progression with an increase in BZ CMs in S-phase compared to M-phase in Pachón (Fig. 5a,b). This fits well with our previously published data, showing that Pachón hearts have similar levels of PCNA-positive BZ cardiomyocytes to surface fish, but these cells fail to multiply ^40^. The main cardiac glucose transporter in *Astyanax*, *slc2a1a*, was strongly expressed in BZ CMs in surface fish, but expression was almost absent in Pachón and BZ expression of glycolysis enzymes *pfkma* and *aldoaa* was reduced (Fig. 5c and Extended Data Fig. 3a). While levels of glycolysis in Pachón hearts were similar to surface fish before injury, they were strongly reduced in Pachón at 7 dpci, and cardiac ATP levels were also lower (Fig. 5d,e). To test if the reduction in glycolysis and resulting low levels of ATP caused the stalling of cell cycle progression, rather than direct clock regulation of the cell cycle, we inhibited glycolysis with 2-Deoxy-d-glucose in surface fish. This indeed recapitulated the halted cell cycle progression in BZ CMs as observed in Pachón and the TKO (Fig. 3a and Fig. 5b,f). Together, these results confirm our findings in the TKO zebrafish that successful cell cycle progression during regeneration is controlled by glycolysis, which in turn is regulated by the core clock mechanism.

**Figure 5:**
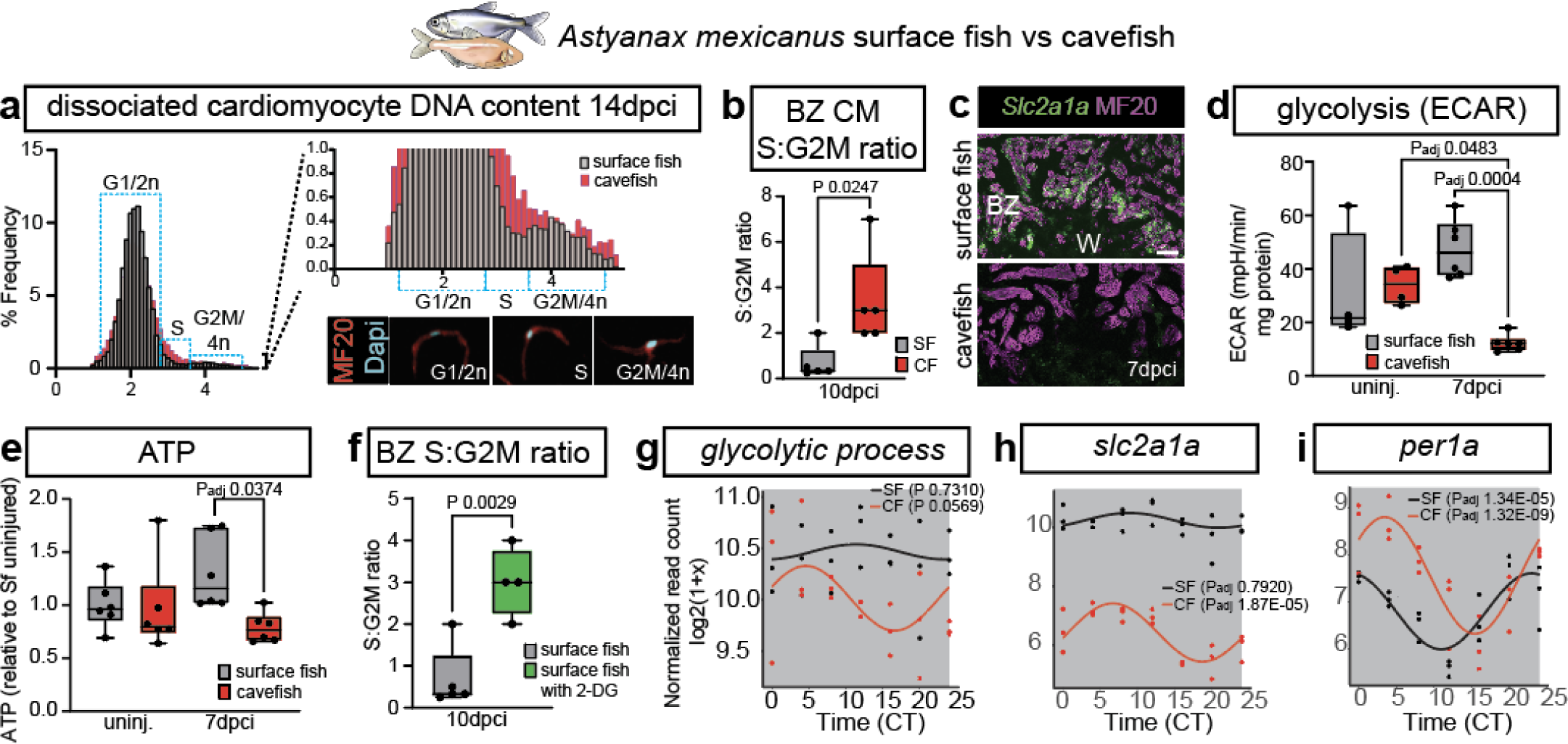
Circadian rhythm and glycolysis regulate cardiomyocyte proliferation during heart regeneration in *Astyanax mexicanus*. **a**, Frequency histogram of DNA intensity of cardiomyocyte nuclei at 14 dpci in Pachón and surface hearts. Images of dissociated cardiomyocytes DNA content, showing G1/2n, S, and G2M/4n nuclei, stained with MF20 and DAPI. **b**, S:G2M ratio comparison of Mef2c+/PCNA+ BZ CM between Pachón and surface at 10 dpci (unpaired two-tailed Student’s t-test). **c**, RNAscope analysis of *slc2a1a* in 7 dpci Pachón and surface heart sections, co-stained with MF20. **d**, **e**, Glycolysis (ECAR) and ATP levels comparison before injury and at 7 dpci between Pachón and surface (one-way ANOVA with Tukey). **f**, S:G2M ratio comparison of Mef2c+/PCNA+ BZ CM between surface treated with 2-Deoxy-d-glucose and control hearts at 10 dpci (unpaired two-tailed Student’s t-test). **g**-**i**, Rhythmic expression of genes representing the *glycolytic process*, *slc2a1a* and *per1a* in Pachón and surface hearts at 7 dpci. (**g** to **i**) *dryR* analysis. Scale bars 100 μm. BZ, border zone cardiomyocytes; W, wound.

Additionally, even though cavefish hearts recapitulated the lower levels of *slc2a1a*, emphasising the role of glycolysis in the resulting stalling of cell cycle progression, the actual amplitude of *slc2a1a* and *glycolysis* rhythm was higher in Pachón (Fig. 5g,h). Indeed, many genes not rhythmic in surface fish specifically gained a rhythm in Pachón, with a clear day/night divide (Fig. 4d). Intriguingly, a day/night pattern was also visible in the genes rhythmic in WT zebrafish that lost their rhythm in the TKO (Extended Data Fig. 2e). Circadian rhythm in cavefish has been shown to be strongly driven by feeding ^54^, suggesting that the absence of light and limited food availability in the cave may have driven circadian regulation to rely more on food entrainment. Consistent with this, core clock genes in Pachón mainly peaked in the morning, corresponding to feeding time, including *per1a* and *per2*, instead of sequential peaking of the different core clock genes as seen in surface fish (Fig. 4b; Fig. 5i and Extended Data Fig. 3b). The phase shift in core clock gene expression in the TKO, which eliminated the morning peak in glycolysis rhythm despite normal feeding and blood glucose levels, alongside the increased glycolysis rhythm and day/night pattern observed in cavefish, made us investigate whether glycolysis itself could be driving this day/night split.

### Insulin resistance inhibits regeneration by resetting the timing of glucose rhythms

During adaptation to cave life, a mutation in the insulin receptor occurred in cavefish, resulting in insulin resistance with hyperglycaemia resembling type 2 diabetes ^55^. Indeed, cardiac glucose levels were higher in Pachón compared to surface fish (Extended Data Fig. 3c). Disruption of the normal rhythm of glucose tolerance is a hallmark of type 2 diabetes ^56^, with a rise in early morning blood glucose levels known as the dawn phenomenon ^57^. This raised the question as to whether the diabetic phenotype, which is not present in the TKO, could reset the disrupted circadian rhythm in cavefish to glucose as the alternative ‘zeitgeber’, a cue that synchronises biological rhythms. The cavefish-specific *insra* mutation has been introduced into zebrafish ^55^, providing an excellent model to specifically test the effect of insulin resistance on glucose rhythms during regeneration. Again, we performed a bulk RNAseq experiment and isolated WT and *insra* mutant ventricles at 4-hour intervals over 24 hours at 7 dpci (Fig. 6a). While the rhythm of most core clock genes was not affected in the *insra* mutant, expression of *per1b* was elevated throughout the 24-hour period, highlighting influence of insulin metabolism on the core clock mechanism (Fig. 6b,c) ^58,59^. Despite the overall normal rhythm of core clock genes, many genes lost their rhythm in the *insra* mutant, including genes involved in insulin signalling (Fig. 6d,e). However, the amplitude of *glycolysis* rhythm was increased (Fig. 6f), similar to that observed in cavefish, highlighting how insulin-resistance creates a unique situation in the fish heart whereby glucose itself may act as a ‘zeitgeber’ in addition to the core clock machinery.

**Figure 6:**
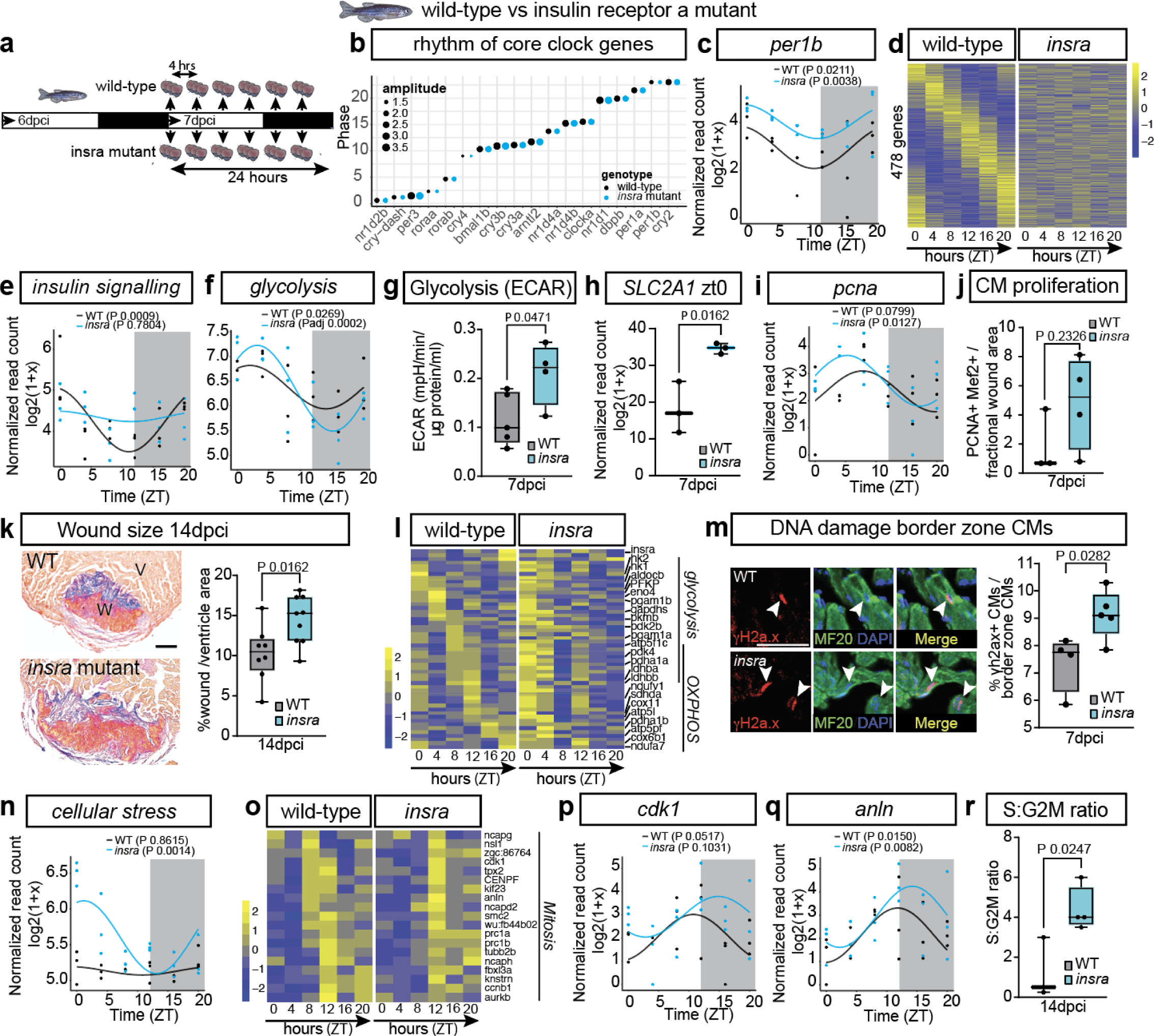
Impact of insulin pathway disruption on heart regeneration through circadian clock-regulated pathways in zebrafish. **a**, RNAseq experimental design schematic. Horizontal bars indicate light (14h) and dark (10h) periods, before and during sampling of 7 dpci WT and *insra* mutant hearts. **b**, Dot plot showing phase and amplitude of core clock genes in WT vs. *insra* hearts at 7 dpci. **c**, Rhythmic expression of *per1b* in WT and *insra* hearts at 7 dpci. **d**, Heatmap of genes identified as rhythmic exclusively in WT 7 dcpi hearts (biological replicates ≥ 3 sampled at 4-hour intervals over 20 hours). **e**,**f**, Rhythmic expression of genes indicative of *insulin signaling* and *glycolysis* between WT and *insra* hearts at 7 dpci. **g**, Glycolysis (ECAR) in WT vs. *insra* hearts at 7 dpci (unpaired two-tailed Student’s t-test). **h**, Expression levels of *SLC2A1* at ZT0 in WT and *insra* hearts at 7dpci (unpaired two-tailed Student’s t-test). **i**, Rhythmic expression of *pcna* in WT and *insra* hearts at 7 dpci. **j**, PCNA/Mef2-positive BZ CM quantification in WT vs. *insra* hearts at 7 dpci. **k**, AFOG staining showing collagen (blue) and fibrin (red) scars in WT vs. *insra* zebrafish hearts at 14 dpci with scar area quantification (unpaired two-tailed Student’s t-test). **l**, Circadian heatmap of g*lycolysis* and OXPHOS subset genes in WT and *insra* at 7 dpci heart. **m**, Immunofluorescence for γH2a.x in MF20+ BZ CMs at 7 dpci (unpaired two-tailed Student’s t-test). **n**, Rhythmic expression of genes representing *cellular stress* in WT and *insra* hearts at 7 dpci. **o**, Circadian heatmap of mitotic subset genes in WT and *insra* hearts at 7 dpci. **p**,**q**, Rhythmic expression of *cdk1* and *anln* in WT and *insra* hearts at 7 dpci. **r**, S:G2M ratio comparison in Mef2c+/PCNA+ BZ CM between WT and *insra* hearts at 14 dpci (unpaired two-tailed Student’s t-test). (**b** to **f**, **i**, **l**, **n** to **q**) *dryR* analysis. Scale bars 100 µm. V, ventricle; W, wound.

While mammalian cardiomyocytes mainly express the insulin-responsive Glut4 transporter (*slc2a4*), fish predominantly rely on Glut1, which facilitates passive glucose transport down a concentration gradient ^60^. As a result, the presence of hyperglycaemia caused by insulin resistance may result in higher cellular uptake of glucose in fish. Indeed, *slc2a1* levels were increased at dawn in the *insra* mutant compared to WT, accompanied by increased glycolysis (Fig. 6g,h). The increase in glycolysis may cause enhanced CM proliferation in *insra* mutants as indicated by the higher peak in *pcna* expression (Fig. 6i,j), however, regeneration was instead negatively affected (Fig. 6k). To better understand the underlying cause, we generated a heatmap of genes involved in cardiac metabolism (Fig. 6l and Extended Data 4a). Strikingly, the metabolic progression seen in WT hearts, from *glycolysis* throughout the light period towards *OXPHOS* at night, was absent in the *insra* mutant. Instead, both *glycolysis* and *OXPHOS* peaked in the morning, recreating the phase shift observed in cavefish (Fig. 5g-i). Both hyperglycaemia and OXPHOS are well-known to cause oxidative stress and DNA damage ^61^, and if OXPHOS is coinciding with DNA replication, this could further exacerbate the damage. There was indeed a strong peak in genes involved in cellular stress in the morning and an accumulation of γh2a.x in *insra* mutant BZ CMs (Fig. 6m,n). Although this did not abolish the rhythmic expression of mitosis-related genes as observed in the TKO and cavefish, it caused a delay in their expression ^46^, including *cdk1* and *anln* (Fig. 6o,q). Nevertheless, similar to loss of rhythm, this delay was associated with impaired cell cycle progression (Fig. 6r). These data confirm the key role that glycolysis rhythms play in cell cycle progression towards mitosis. Under normal conditions, metabolic rhythms progress diurnally, but circadian misalignment disrupts this pattern. In insulin resistant, however, high glucose levels after feeding can re-synchronise these rhythms to peak in the early morning. Therefore, successful cell cycle progression of cardiomyocytes during regeneration requires a tight circadian regulation of glucose metabolism with both the internal biological clock and glucose as interconnected zeitgebers.

## Discussion

Until now, the influence of circadian rhythms on heart regeneration remained unknown. Our study highlights the critical role of a balanced diurnal rhythm in glucose metabolism for successful cardiomyocyte proliferation and regeneration, with both the circadian clock and glucose signalling regulating this rhythm. A glycolysis rhythm that lacks amplitude leads to a complete lack of upregulation of genes involved in mitosis, while an excessively high rhythm delays mitosis, similarly preventing cells from progressing into M-phase due to DNA damage. During successful regeneration, gene expression of key processes peaks at distinct time points over 24 hours, indicating that the cardiomyocyte cell cycle indeed takes 24 hours to complete ^62^ with unique cellular behaviour depending on the time of sampling.

In particular, genes involved in mitosis become highly rhythmic after injury and this rhythm is lost in both the TKO and cavefish. As core clock genes can directly control cell cycle regulators ^63–65^, this loss might partially be due to loss of direct regulation of the cell cycle by core clock genes. However, our data shows that inhibiting glycolysis is sufficient to recapitulate the cell cycle stalling. While glycolysis is known to be crucial for proliferation of border-zone cardiomyocytes ^48^, our data pinpoints its specific role in progression to M-phase. Insufficient energy to complete DNA replication and support DNA repair mechanisms aligns with the increased γH2a.x staining in TKO BZ CMs and upregulation of checkpoint genes, indicating that DNA damage prevents progression into M-phase.

Glycolysis peaks early morning, around feeding time. Circadian misalignment in the TKO mutant, due to a phase advance of all main regulators of the core clock system, leads to a loss of *slc2a1* rhythm, indicating lack of glucose uptake with disruption of downstream glycolysis, despite normal feeding and blood glucose levels. This suggests that regulation of *SLC2A1* is directly clock-controlled, similar to *SLC2A4* (Glut4), the main glucose transporter in mammalian cardiomyocytes, which is known to have E-box sites in the promoter ^66^. An E-box is a *cis*-element required for circadian transcription by clock genes ^67^. Similar to *SLC2A4*, we find a consensus E-box element (TTACGTAA) in the promoter of *Astyanax Slc2a1a* and a non-canonical E-box element in the zebrafish *slc2a1* promoter (Extended Data Fig. 4b). Therefore, misalignment between circadian clock expression and feeding can explain why *slc2a1* cannot be activated in the morning when needed in the TKO due to a core clock phase advance, as well as the alignment to altered delayed core clock genes expression in cavefish. While advances or delays in rhythms influence metabolic processes in distinct ways ^44^, we show that both result in reduced regeneration, highlighting the requirement of a balanced glucose rhythm for successful regeneration. The presence of a day-night rhythm in the WT, as highlighted by its loss in the TKO, as well as sequential 24-hour rhythm progression, shows that both light and food are zeitgebers in the WT hearts, emphasising the complex interplay between these processes ^54^.

Misalignment of circadian rhythm and timing of meals is a key factor in the development of diabetes ^68–70^, as glucose tolerance varies during the 24-hour period and mouse models lacking core clock genes show a diabetic phenotype ^59,71^. While cavefish indeed have hyperglycaemia, this will also be caused by the mutated insulin receptor ^55^ and possibly other specific clock and metabolic related mutations that occurred during evolution in the cave environment, including mutations in *per2* and *MC4R* ^72–74^. The combination of several factors driving hyperglycaemia is likely crucial to completely reset cavefish circadian rhythm to food and glucose as alternative zeitgebers, as this is not seen in the *insra* zebrafish mutant. While it is known that insulin can reset the clock via regulation of PER ^75^, replication of the cavefish *insra* mutation showed that insulin resistance in fish can increase *per1b* levels but does not completely reset the clock mechanism. The fish heart shows a counterintuitive response to insulin resistance, with enhanced glycolysis at the time of feeding. While this could theoretically enhance proliferation, it also results in high levels of OXPHOS and oxidative stress at the time of DNA replication. The peak in OXPHOS is normally during the night, temporally separated from DNA replication, avoiding DNA damage caused by generation of ROS ^76^.

Future research on heart regeneration should account for the influence of circadian rhythms as they directly regulate the processes crucial for successful regeneration in a temporal manner. Understanding how misalignment between feeding and the circadian clock affects cardiomyocyte function across different species, including humans, could lead to breakthroughs in addressing heart regeneration. Chronotherapy or pharmacological modulation of clock genes or their downstream targets could be a potential strategy to synchronise metabolic rhythms and promote effective heart repair. Our study is the first to investigate how circadian rhythms influence heart regeneration, specifically through glucose metabolism and cardiomyocyte cell cycle progression. However, their impact likely extends further, influencing processes such as extracellular matrix deposition, which governs scar formation and resorption, and represents an important area for future study.

## Methods

### Animals

Zebrafish and *Astyanax mexicanus* were used in this study. The *cry1a*, *cry3a* and *per2* mutant (referred to as TKO, triple knock-out) ^43^ and wild-type controls were kept on TL background. The homozygous P211L *insra* mutant ^55^ and control zebrafish lines were kept on KCL background. All zebrafish were kept at 28°C under a 10/14- or 12/12-hours light/dark cycle. *Astyanax mexicanus* surface and Pachón cavefish were obtained from Yoshiyuki Yamamoto at UCL, bred for 1-3 generations at Oxford and maintained at 20-21°C under a light/dark cycle of 12/12 hours. All procedures involving animals at the University of Oxford were approved by the local animal experiment committees and performed in compliance with animal welfare laws, guidelines, and policies according to national law. All experiments involving zebrafish at Ulm University were approved by the state of Baden-Württemberg and the animal care representatives of Ulm University.

### Light conditions

Animals were maintained on a 12/12 or 14/10 light-dark (LD) cycle prior to the experiment. During LD, time was expressed as Zeitgeber Time (ZT), with ZT0 defined as the time of lights-on and ZT12 or ZT14 as lights-off. After transfer to constant darkness (DD), time was expressed as Circadian Time (CT), with CT12 defined as the projected onset of activity, corresponding to the animal’s subjective dusk.

### Cardiac cryoinjury

Zebrafish and *Astyanax mexicanus* aged ∼1-1.5 year were used. Cryoinjury of the ventricle was performed per previous description. In brief, fish were anaesthetised in MS-222 (250 mg/L, Sigma-Aldrich) and placed ventral side facing upwards in a sponge-holder under a dissection microscope (Olympus). An incision was made using forceps and microdissection scissors at the level of the heart, which facilitated the exposure of the ventricle out of the pericardial cavity. Cryoinjury was induced on the tip of the ventricle using a copper probe pre-chilled in liquid nitrogen. Following surgery, fish were returned to fresh tank water immediately and their gills were pipetted through using water to facilitate breathing and restoring of swimming capability. Hearts were isolated at the indicated timepoints and processed for further analysis.

### Sample processing for histology and immunofluorescent staining

Isolated hearts were rinsed in pre-cooled PBS and fixed in 4% paraformaldehyde (PFA, ChemCruz) overnight at 4°C. Dehydration was performed using increasing ethanol concentrations (70%, 80%, 90%, 96% and 100%). After overnight incubation in 100% 1-butanol, the samples were placed in paraffin (Paraplast, Sigma) at 65°C and sectioned using a Microm HM325 microtome at 7-10 µm thickness (or 25 µm for cell cycle analysis). The sections were mounted on SuperFrost glass slides (VWR) and dried overnight at 37°C for further processing.

### Acid Fuchsin Orange G (AFOG) staining

Following heart sectioning, one out of every ten sections was mounted per sample to ensure comprehensive coverage for AFOG staining of the entire heart. The procedure was performed as previously described ^40^. In summary, the samples underwent dewaxing twice for 6 mins each in Histo-Clear II (National Diagnostics) and were subsequently rehydrated in a descending concentration of ethanol (100%, 96%, 90%, 80%, 70%, 1 min each). The specimens were fixed in Bouin’s solution (VWR) at 60°C for two hours and then treated with Phosphomolybdic acid, 1% solution for 5 min. Finally, the specimens were stained in the AFOG solution (0.5% w/v methyl blue, 1% w/v orange G, 1.5% w/v acid fuchsin, pH 1.09) for 7 mins. The stained slides were dehydrated in ascending concentrations of ethanol (70%, 80%, 90%, 96%, 100%, 1 min each) and washed twice for 6-min in Histo-Clear II (National Diagnostics). Finally, the slides were briefly allowed to dry before being mounted in dibutylphthalate polystyrene xylene (DPX) medium (Sigma-Aldrich) in preparation for imaging.

### Immunofluorescent staining

For the immunohistochemistry analysis, a minimum of three sections per heart, featuring clear wounds, were selected for subsequent processing. The samples underwent dewaxing with xylene and rehydration as previously described. The sections were boiled in antigen unmasking solution, citric acid-based (Vector Laboratories), for 8 mins using a pressure cooker, followed by a 20-min cooldown period in PBS. To mitigate non-specific binding, sections were incubated in TNB blocking buffer for 30-60 mins at room temperature. Incubation with the primary antibody solution occurred overnight in a humidified chamber at room temperature. Following a wash with PBST (containing 0.1% Tween-20), the sections underwent a 2-hour incubation with secondary antibodies at room temperature. Primary antibodies, Mef2c (Biorbyt Cat# orb1336562), MF20 (DSHB), PCNA (Clone PC10, Dako), BrdU (Abcam Cat# ab6326), and secondary antibodies, Alexa Fluor® 488 and Alexa Fluor® 568 (Invitrogen), were prepared in TNB buffer at a ratio of 1:200. Upon completion of staining, the secions were mounted in ProLong Gold Antifade Mountant (Invitrogen) in preparation for imaging.

For γh2a.x staining cryoinjured hearts were fixed in 4% PFA for 2 h at RT. Samples were then washed three times for 10 minutes in 4% sucrose/phosphate buffer and incubated overnight at 4 °C in 30% sucrose/phosphate buffer for cryoprotection. Hearts were embedded in NEG-50 frozen section medium (ThermoFisher, Cat#1214849) using Peel-A-Way Embedding Molds (Sigma, Cat#18646A-1), and cryosectioned at a thickness of 10 μm. Sections were evenly distributed across six serial slides to ensure uniform representation of all ventricular areas.

The slides were washed three times with PEMTx buffer (80 mM Na-PIPES, 5 mM EGTA, 1 mM MgCl₂, pH 7.4, 0.2% Triton X-100), followed by a single wash with PEMTx supplemented with 50 mM NH₄Cl. Blocking was performed in PEMTx/NGS buffer (10% Normal Goat Serum, 1% DMSO, 89% PEMTx) for 1 hour at room temperature in a humidified chamber. Primary antibodies, histone H2A.XS139ph (phospho Ser139) (Genetex Cat# GTX127342, 1:2000), Mf20 (DSHB, Cat# MF20, 1:100) were applied in PEMTx/NGS and incubated overnight at 4 °C. Secondary antibodies Alexa Fluor™ 555 and Alexa Fluor™ 488 (Invitrogen Cat# A21429 and Cat# A11034) were used at a dilution of 1:1000. Nuclei were shown by DAPI (40,60-diamidino-2-phenylindole) staining. Slides were mounted with Fluorsave (Merck Cat #345789) mounting medium.

### RNAscope

Paraffin sections were first mounted on SuperFrost slides (VWR) and dried overnight at 37°C. These were then incubated for 1 hour at 60°C and then deparaffinized with two 5 mins xylene washes before two washes in 100% ethanol. Sections were then left to air dry, which was followed by treatment with 3% hydrogen peroxide for 10 mins at room temperature. Next, sections were boiled (98-102°C) with RNAscope target retrieval for 15 mins, which was followed by incubation with RNAscope protease III at 40°C for 12 mins. Following protease treatment, sections were incubated with probes for 2 hours at 40°C. To detect the hybridization signal, RNAscope Multiplex Fluorescent Detection Reagents v2 (ACD, 323100) utilising the TSA Plus Cyanine 3 or Cyanine 5 fluorophore (Perkin Elmer, NEL744001KT) were applied according to the manufacturer’s instructions. RNAscope was imaged on an Olympus FluoView3000 confocal microscope.

### Bulk RNA sequencing and data analysis

Total RNA was extracted from samples using the Quick-RNA Microprep Kit (Zymo Research, Cat. No. R1050) according to the manufacturer’s instructions. This kit enables rapid purification of high-quality RNA from small sample volumes, with on-column DNase treatment to eliminate genomic DNA contamination. RNA yield and purity were initially assessed using the Qubit 3.0 Fluorometer (Thermo Fisher Scientific) with the RNA HS Assay Kit (Thermo Fisher Scientific). RNA integrity was further evaluated using the Agilent 4200 TapeStation System with RNA ScreenTape (Agilent), providing RNA Integrity Number equivalent (RINe) scores to ensure sample suitability for downstream applications. Library preparation was performed using QuantSeq 3’ mRNA-Seq V2 Library Prep Kit FWD with UDI 12 nt Set B1, (UDI12B_0001-0096, 96 preps). The optimal number of cycles for the endpoint PCR of the QuantSeq V2 was determined using 192.96 kit + 020 (PCR Add-on Kit for Illumina, 96 rxn) according to the manufacturer’s instructions. Following library construction, fragment size distribution and library quality were confirmed using the Agilent 4200 TapeStation System with D1000 ScreenTape (Agilent). Libraries with appropriate concentration and size distribution were pooled in equimolar amounts for sequencing. The libraries were sequenced on Illumina next-generation sequencing (NGS) platforms as a service provided by Genewiz (Azenta Life Sciences) and the Wellcome Centre for Human Genetics (WCHG) sequencing facilities.

RNA sequencing reads were trimmed of adapters and checked for quality with Trim Galore! (v.0.6.7) ^77^ before alignment and gene counting using STAR (v.2.7.10a ^78^) against the zebrafish genome (GRCz11) with Ensembl (https://ensembl.org) annotations (release 110). For *Astyanax* samples, alignment was against the Mexican tetra (v.2.0) genome with customised Ensembl annotations (release 103). Gene counts were loaded in R (v.4.0.1) ^79^ and genes with no counts in any sample were removed before further analysis with the R package *dryR* (v.1.0.0) ^41,80^ to identify rhythmic genes. Rhythmicity of pathways was evaluated by first constructing a pathway metagene by averaging the counts of genes comprising the pathway for each sample and adding the metagene to the raw counts matrix before running *dryR*. Heat maps were generated using the R package pheatmap ^81^ using a script adapted from *dryR* “plot_models_rythm” to arrange genes in ZT order based on their phase. Where two groups are shown separated by a gap (e.g., wild-type and TKO) in the same heat map, genes (rows) are in the same order for both groups and mean values per timepoint are scaled across all samples. Dot plots were created using the R package ggplot2 (v.3.4.3) ^82^ with phase and amplitude from *dryR* analyses.

### RNA extraction and RT-qPCR analysis

Total RNA was extracted from samples using the Quick-RNA Microprep Kit (Zymo Research, Cat. No. R1050) according to the manufacturer’s instructions. RNA yield and purity were initially assessed using the Qubit 3.0 Fluorometer (Thermo Fisher Scientific) with the RNA HS Assay Kit (Thermo Fisher Scientific). Reverse transcription was performed using the High-Capacity cDNA Reverse Transcription Kit (Thermo Fisher Scientific) according to the manufacturer’s protocol. The synthesized cDNA was stored at −20°C until further use. Quantitative real-time PCR was performed using the QuantStudio 5 platform (Applied Biosystems), with Fast SYBR™ Green Master Mix (Thermo Fisher Scientific). The cycling conditions were as follows: initial denaturation at 95°C for 20 seconds, followed by 40 cycles of denaturation at 95°C for 3 seconds, annealing/extension at 60°C for 30 seconds. The total runtime for RT-qPCR was approximately 45 minutes. Melt curve analysis was conducted to confirm the specificity of the amplified products. Primers were designed using Primer-BLAST and synthesized by Integrated DNA Technologies. Primer specificity was verified by BLAST analysis and gel electrophoresis of PCR products. Primer efficiency was determined by generating a standard curve using serial dilutions of cDNA, and only primers with efficiencies between 90-110% (R² > 0.98) were used. Normalisation and Data Analysis Gene expression levels were normalised against the housekeeping gene(s). Relative expression was calculated using the 2^(−ΔΔCt) method. All reactions were performed in technical triplicates. All data were expressed as mean ± standard deviation (SD) or standard error of the mean (SEM). Differences between groups were assessed using one-way ANOVA in GraphPad Prism. A p-value < 0.05 was considered statistically significant.

### Seahorse metabolic assays

Zebrafish and *Astyanax* ventricles were isolated and deposited in separate wells of an XFe24 Islet Capture Microplate (Agilent) in Seahorse XF DMEM medium, pH 7.4 (supplemented with 2 mM glutamine, 1 mM pyruvate, and 10 mM glucose without phenol red). Measurements were performed with the XFe24 analyser (Agilent) at room temperature. For *Astyanax mexicanus,* extracellular acidification rate (ECAR), Seahorse XF Cell Energy Phenotype Test Kit was used which employs a simultaneous injection of oligomycin (2.5 μM) and FCCP (1 μM) to create stressed conditions in terms of energy production. For zebrafish ECAR, the XF Cell Mitostress Test Kit (Agilent) was used which employs serial injections of oligomycin (50μM), FCCP (30μM) and rotenone/antimycin A (45μM) as previously described (Lekkos et al., unpublished). The data presented is a result of the average of the top three FCCP measurements. Following the Seahorse assays, protein was isolated from the hearts and protein concentration was measured using BCA Protein Assay (Pierce) to normalise metabolic measurements.

### Glucose and ATP measurements

For cardiac glucose and ATP measurements, *Astyanax* ventricles were isolated and lysed using the Fluorometric Glucose Assay Kit (Abcam, ab6533) or Fluorometric ATP Assay Kit (Abcam, ab83355), according to manufacturer’s instructions. Prior to the recommended deproteinisation step, protein was isolated from the heart lysates and protein concentration was measured using BCA Protein Assay (Pierce) to normalise glucose or ATP levels. Deproteinisation was performed using Deproteinising Sample Preparation Kit – TCA (Abcam, ab204708). For blood glucose measurements, whole blood was collected from zebrafish and glucose levels assessed using the Accu-Chek® Instant Blood Glucose System, according to manufacturer’s instructions.

### Drug treatment

*Astyanax* surface fish received intraperitoneal injections once daily with either a DMSO control or 2-deoxyglucose (Sigma-Aldrich, 1 mg/g) for five consecutive days, from 5 to 9 dpci and hearts isolated at 10 dpci. Zebrafish WTs and TKO mutants received one intraperitoneal injection with BrdU (Invitrogen) at 7 dpci and hearts were isolated at 14 dpci. The drug solutions were prepared in 0.01 µM DMSO (Sigma-Aldrich) in saline and injected using a BD Micro-fine U100 insulin Syringe (gauge 30G) with injection volume of 20 µL.

### Imaging

RNAscope and immunohistochemistry images of heart samples were acquired using a Nikon, Eclipse Ci-L microscope with Nikon DS-Fi3 camera (Nikon, Tokyo, Japan), Zeiss LSM 880 and FluoView 3000 Olympus confocal laser scanning microscopes.

### Image analysis

For AFOG analysis, we quantified wound size using Fiji ImageJ software on minimum three sections per heart with the largest wound or on sections equally covering the whole ventricle volume. The wound size was presented as a percentage of wound area against the ventricular area or as a percentage of wound volume against the ventricle volume. For proliferation measurements, we counted Mef2c+/ PCNA+ double-labelled nuclei and those labelled only by Mef2c+ within the border zone (BZ), a 100 µm region adjacent to the wound border. The percentage of proliferating cardiomyocytes was obtained by dividing the number of PCNA+/Mef2c+ against the total Mef2c+ nuclei.

For cell cycle analysis using paraffin sections, hearts were sectioned at 25 µm to capture whole nuclei and stained with DAPI, PCNA, and Mef2c antibody (or BrdU and MF20 for the TKO experiment in Fig.3). The images were taken with a Zeiss 880 or FluoView 3000 Olympus confocal microscope using the same setting of laser power, voltage, offset, and pinhole across the board. Z-stack images were taken with 0.9 µm for each step, with approximately 25 – 30 steps per section. Cardiomyocyte DNA content was measured in z-stacks using Imaris Version 8.2 software (Bitplane AG, Switzerland). Nuclei were identified and segmented using the “Surfaces” tool based on PCNA or BrdU signal, then cardiomyocytes were selected based on Mef2c (nuclear cardiomyocyte marker) or MF20 (cytoplasmic cardiomyocyte marker) signal. The same “Surfaces” parameters for nuclei identification were used across all images. Integrated intensity of the DAPI signal was measured in every identified nucleus and exported for further analysis.

For cell cycle analysis using cardiomyocyte suspensions, DNA content was determined using a pipeline made in Cell profiler, which was applied to each field of view; the number of nuclei within each cell was also recorded. In brief, images firstly underwent illumination correction to correct for any inconsistent illumination across the image. Next, using the “IdentifyPrimaryObjects” module, nuclei were thresholded based on the DAPI signal and selected as the primary objects. This was followed by using the “IdentifySecondaryObjects” module to identify and select cardiomyocytes based on the MF20 signal. The “IdentifySecondaryObjects” module was set to threshold on MF20 signal by propagating from identified nuclei. Next, cardiomyocyte and non-cardiomyocyte nuclei, were identified using the “MaskObjects” module, which masked identified nuclei (primary objects) by the cytoplasmic MF20 signal (secondary objects). Lastly, Using the “MeasureObjectIntensity” module the integrated intensity of the DAPI signal was measured in every identified nucleus and exported for further analysis.

For both cell cycle analysis approaches described above, intensity measurements between images were normalised by performing the below calculation on each nucleus:

*DNA content* = *Integrated Intensity of nucleus x 2 Median Integrated Intensity of all nuclei in image*

Ultimately, the resulting value represents the DNA content of a given cell. DNA content measurements of all cardiomyocyte nuclei across every heart analysed (both surface fish and Pachón) were used to generate a frequency histogram using a bin size of 0.1n in GraphPad Prism 9. Based on the histogram peaks of surface fish cardiomyocytes, the phase or ploidy state was estimated for each surface and Pachón cardiomyocyte.

For γh2a.x analysis, images were acquired as single optical planes using 20 x magnification with a Leica Sp8 confocal microscope. Quantification was performed manually on 3 sections per heart that contained the largest wounds. Analysis was limited to myocardium within 150 µm of the wound border. The wound area was defined using immunofluorescence for the cardiomyocyte markers Mf20. In ImageJ, the wound border was outlined with the “freehand selection” tool and saved to the Region of Interest (ROI) manager. The selection was then expanded by 150 µm and added as a second ROI. Both ROIs were selected, and the XOR function with overlays was applied to define the final analysis region.

### Heart dissociation

To generate single cell suspensions, *Astyanax* hearts were isolated at 14 dpci after cryoinjury into ice-cold 1xPBS. Next, the atrium and the bulbous arteriosus were removed before bisecting the ventricle in two. Tissue was then washed twice in ice cold dissection buffer (3% BSA, 20 mM glucose in 1xPBS) and incubated in a solution containing 0.2% trypsin, 0.8 mM EDTA (Gibco) supplemented with 20 mM glucose (Sigma) and 10 mM 2,3-butanedione monoxime (BDM, Sigma) at 4°C with gentle agitation. Following trypsin incubation, ventricles were washed three times in dissection buffer supplemented with 10 mM BDM and transferred to a 24-well culture plate, where ventricles were digested in Accumax (EMD Millipore) supplemented with 20 mM glucose and 10 mM BDM (digestion buffer) for 15 mins at room temperature under mild agitation. After 15 mins, the digestion buffer was removed and added to an equal volume of stopping buffer (30 mM Taurine, 5.5 mM Glucose, 10 mM 2,3-Butanedione Monoxime, 10 mM HEPES 12.5 µM CaCl_2_, 10% FBS in 1x PBS) to neutralise the digestion. Simultaneously, undigested tissue fragments were transferred to a new well with fresh digestion buffer and gently pipetted up and down before incubating under mild agitation at room temperature. This was repeated until the tissue fragments had fully digested. After digestion, dissociated cells were pelleted by centrifugation at 400 g for 5 mins before being resuspended in 2% PFA for 30 mins on ice (Santa Cruz, sc-281692). Following fixation, cells were again centrifuged at 400 g for 5 mins and resuspended in 1xPBS. Cells were spread on Superfrost Plus slides (ThermoFisher Scientific) and air dried overnight.

### Statistics

Data was plotted using GraphPad Prism V10.1.1 (GraphPad Software Inc., USA) and presented as mean ± SD or SEM. Data were analysed using unpaired two-tailed Student’s t-test, one-way ANOVA with Tukey’s multiple comparisons test, or two-way ANOVA with Sidak’s multiple comparisons test. The statistical test and animal numbers used in each experiment were included in the figures or their legends. Differential rhythmicity analysis was performed using *dryR, a* statistical framework based on model selection that is developed to identify and estimate changes in rhythmic parameters (amplitude and phase) and mean expression in various conditions.

## Acknowledgments

We would like to thank Dr. Jun Hirayama for sharing the TKO, Prof. Nicolas Rohner for providing the *insra* mutant and Prof. Paul Riley for critically reading the manuscript. We would also like to thank the Animal Facility for their care for the animals.

## Funding

European Research Council (ERC) under the European Union’s Horizon 2020 research and innovation program 715895, CAVEHEART, ERC-2016-STG (MTMM)

British Heart Foundation project grant PG/15/111/31939 (MTMM)

British Heart Foundation project grant PG/23/11189 (GR, MTMM)

BHF Centre of Research Excellence Oxford (RE/13/1/30181) (MTMM)

King Faisal Specialist Hospital & Research Centre Post-doctoral fellowship (RA)

Oxford Medical Research Council Doctoral Training Program MR/W006731/1 (KL)

DFG (German Research Foundation) CRC1506 project-ID 450627322 (GW)

DFG (German Research Foundation) CRC1149 project-ID 251293561 (GW)

DFG (German Research Foundation) CRC1279 project ID 316249678 (GW)

## Author contributions

Conceptualization: GR, MTMM

Methodology: GR, RA, WTS, DPP, KL, JWT, MEL, MTMM

Investigation: GR, RA, WTS, DPP, KL, SA, VO, MEL, MTMM

Visualization: GR, RA, WTS, MEL, MTMM

Funding acquisition: GR, RA, GW, MTMM

Project administration: MTMM

Supervision: GW, MTMM

Writing – original draft: GR, RA, MTMM

Writing – review & editing: GR, RA, GW, MTMM

## Extended Data Figures

**Extended Data Fig. 1.**
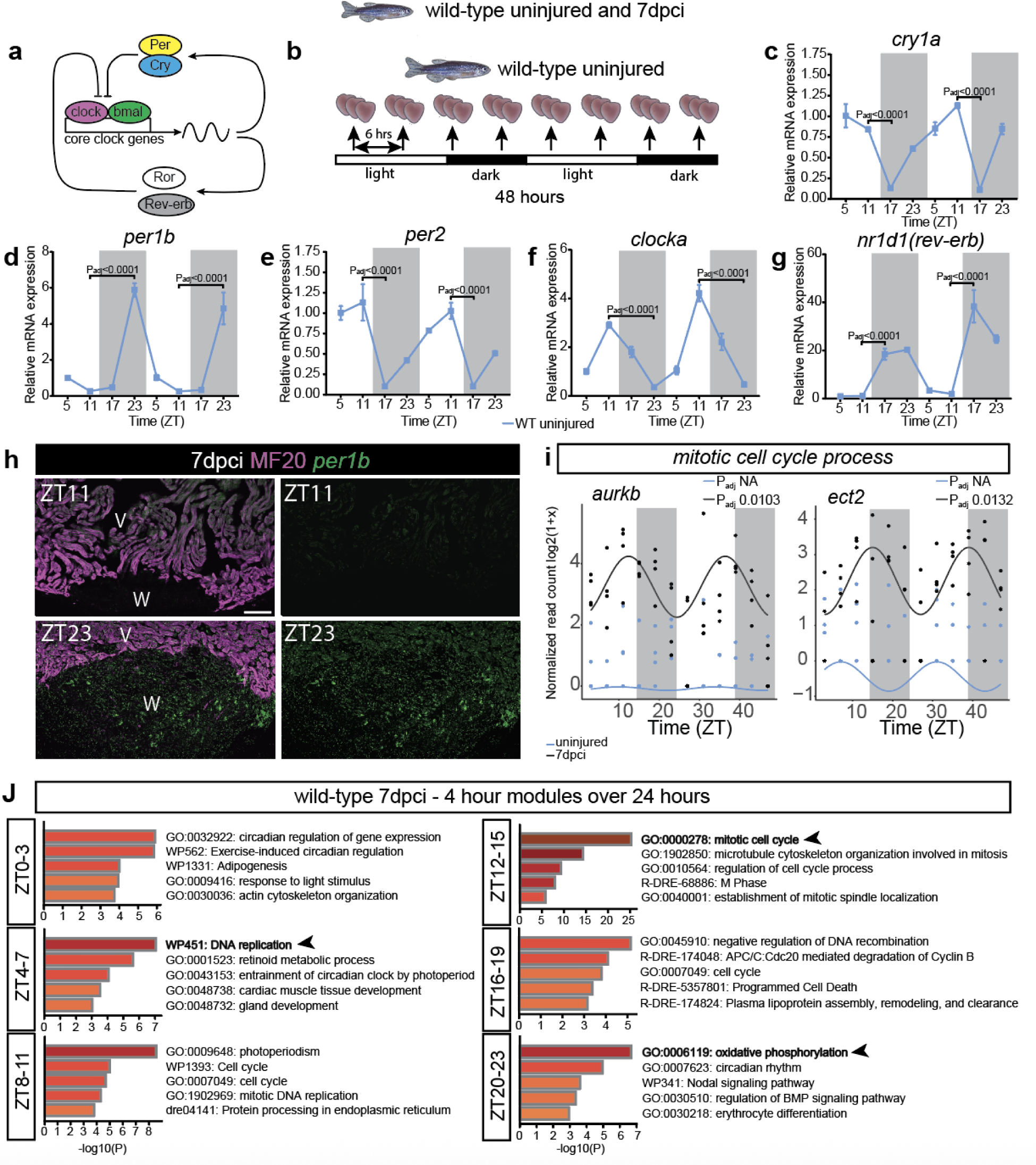
Characterisation of the circadian clock gene expression in WT zebrafish hearts before and after cryoinjury. **a**, Simplified schematic of the core clock mechanism (Transcription/Translation feedback loop). **b**, RT-qPCR experimental design schematic. Horizontal bars indicate light (14h) and dark (10h) periods before and during sampling of uninjured zebrafish WT hearts. **c-g**, RT-qPCR analysis of the rhythmic expression of *cry1a*, *per1b*, *per2* and *nr1d1* in uninjured zebrafish WT hearts (one-way ANOVA with Tukey). **h**, RNAscope analysis *per1b* in 7 dpci WT zebrafish heart sections, co-stained with MF20. **i**, Rhythmic expression profiles of *aurka* and *ect2* in uninjured vs. 7 dpci WT zebrafish hearts. **j**, Bar graph of Metascape-enriched terms for CCGs in WT zebrafish hearts at 7 dpci, coloured by p-values. The analysis was performed on 4-hour time windows over 24h, with each module including a subset of genes peaking at the specific time window.

**Extended Data Fig. 2.**
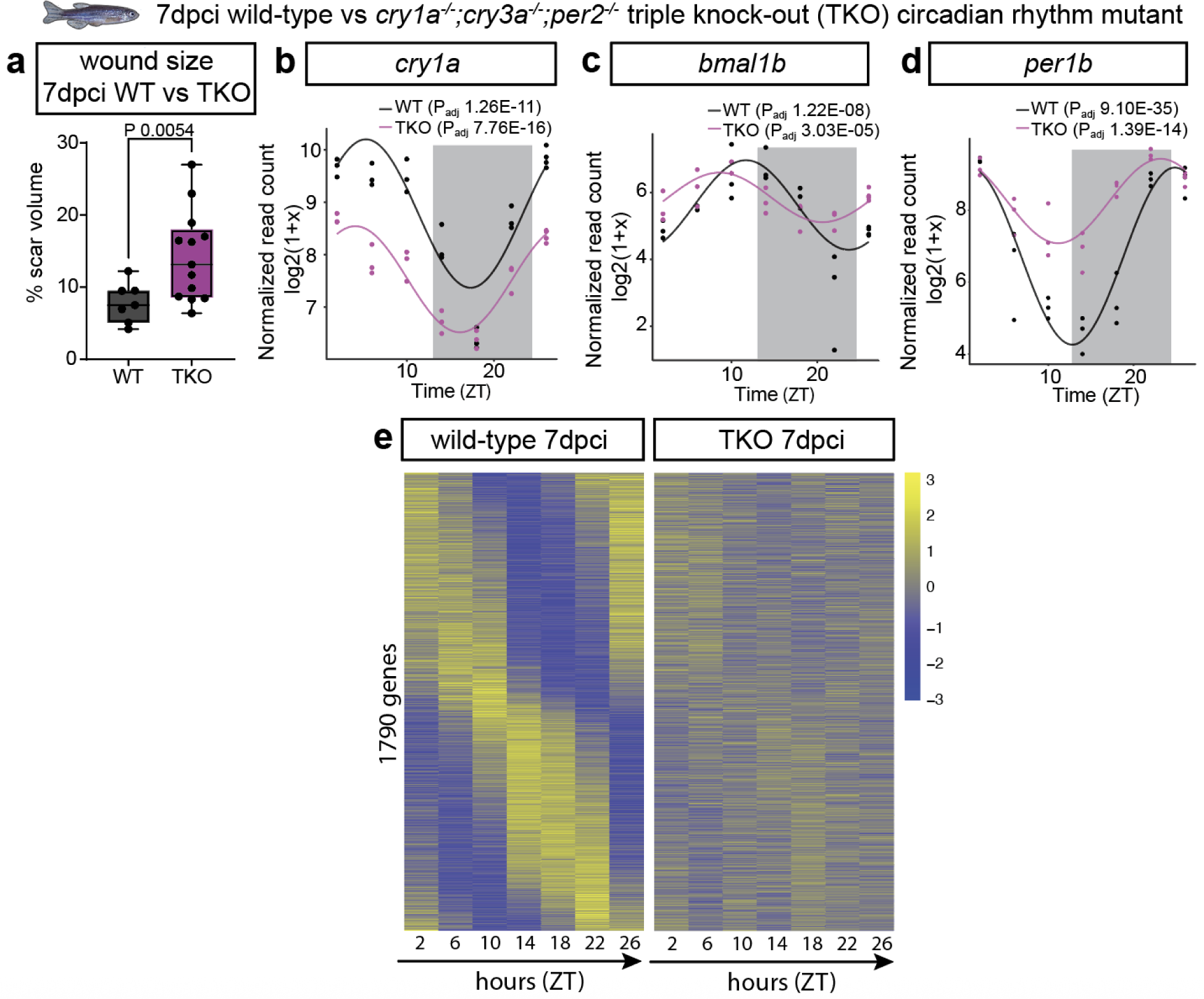
Analysis of the rhythmic expression and regenerative capability between WT and TKO zebrafish hearts. **a**, Scar volume quantification in WT vs TKO zebrafish hearts at 7 dpci (unpaired two-tailed Welch t-test). **b**-**d,** Rhythmic expression profiles of *cry1a*, *bmal1b* and *per1b* in WT vs TKO zebrafish hearts at 7 dcpi. **e**, Heatmap of genes identified as exclusively rhythmic in WT hearts at 7 dpci using *dryR*.

**Extended Data Fig. 3.**
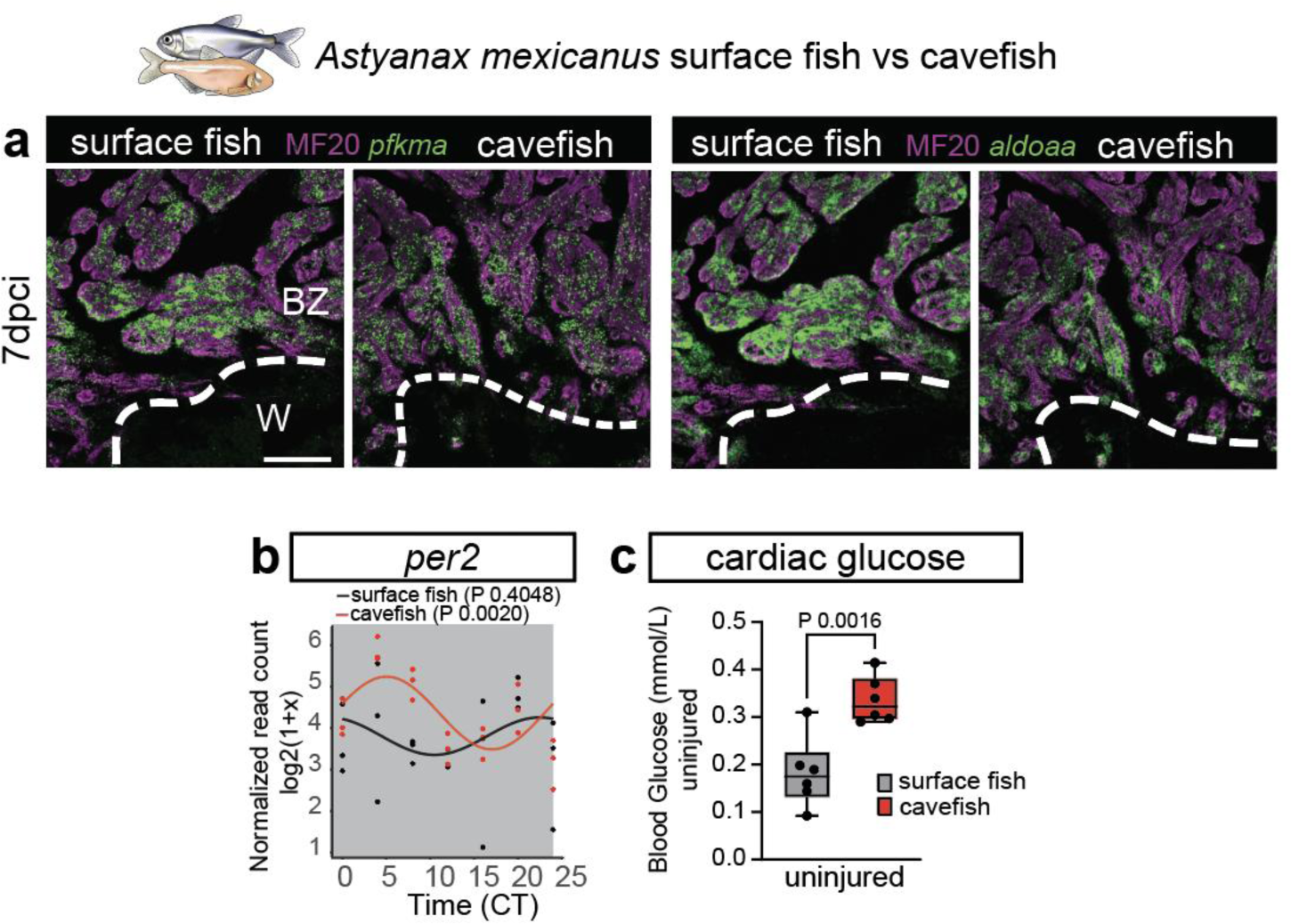
Glycolytic metabolism and *per2* rhythmic expression during heart regeneration in *Astyanax Mexicanus*. **a**, RNAscope analysis of *pfkma* and *aldoaa* in 7 dpci Pachón and surface heart sections, co-stained with MF20. **b**, Rhythmic expression profile of *per2* in Pachón and surface fish hearts at 7 dpci. **c**, Cardiac glucose quantification in uninjured hearts between Pachón and surface fish (unpaired two-tailed Student’s t-test).

**Extended Data Fig. 4.**
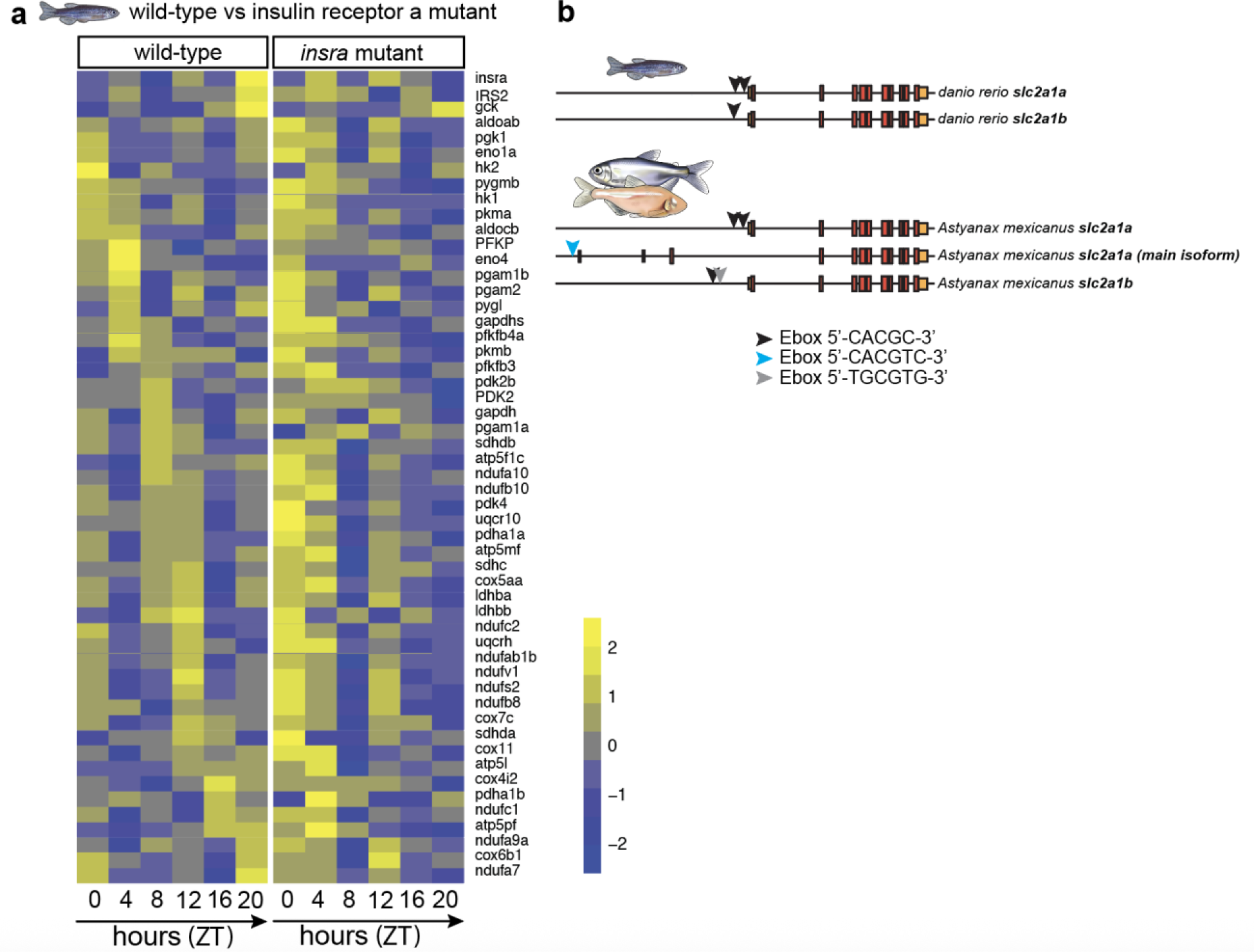
Analysis of the glycolysis and OXPHOS rhythmic expression in WT versus *insra* mutant hearts. **a**, Circadian heatmap of the complete list of g*lycolysis* and *OXPHOS* subset genes in WT and *insra* mutant hearts at 7 dpci. **b**, Schematic of presence of E-box elements in the promoters of *slc2a1a* and *slc2a1b* in zebrafish and *Astyanax mexicanus*.

